# Characterization of methionine dependence in melanoma cells

**DOI:** 10.1101/2023.04.05.535723

**Authors:** Sarita Garg, Lauren C. Morehead, Jordan T. Bird, Stefan Graw, Allen Gies, Aaron J. Storey, Alan J. Tackett, Rick D. Edmondson, Samuel G. Mackintosh, Stephanie D. Byrum, Isabelle R. Miousse

**Affiliations:** Department of Biochemistry and Molecular Biology, University of Arkansas for Medical Sciences

**Keywords:** Methionine, methylation, proteomics, melanoma, cell cycle, ribosome

## Abstract

Dietary methionine restriction is associated with a reduction in tumor growth in preclinical studies and an increase in lifespan in animal models. The mechanism by which methionine restriction inhibits tumor growth while sparing normal cells is incompletely understood. We do know that normal cells can utilize methionine or homocysteine interchangeably (methionine independence) while most cancer cells are strictly dependent on methionine availability. Here, we compared a typical methionine dependent and a rare methionine independent melanoma cell line. We show that replacing methionine, a methyl donor, with its precursor homocysteine generally induced hypomethylation in gene promoters. This decrease was similar in methionine dependent and methionine independent cells. There was only a low level of pathway enrichment, suggesting that the hypomethylation is generalized rather than gene specific. Whole proteome and transcriptome were also analyzed. This analysis revealed that contrarily to the effect on methylation, the replacement of methionine with homocysteine had a much greater effect on the transcriptome and proteome of methionine dependent cells than methionine independent cells. Interestingly, methionine adenosyltransferase 2A (MAT2A), responsible for the synthesis of s-adenosylmethionine from methionine, was equally strongly upregulated in both cell lines. This suggests that the absence of methionine is equally detected but triggers different outcomes in methionine dependent versus independent cells. Our analysis reveals the importance of cell cycle control, DNA damage repair, translation, nutrient sensing, oxidative stress and immune functions in the cellular response to methionine stress in melanoma.

## Introduction

Fundamental differences in metabolism between normal and cancer cells represent optimal targets for cancer therapy, minimizing adverse effects. One such fundamental difference was identified over 40 years ago in the way normal cells and cancer cells respond to methionine depletion, and more particularly in their response to methionine depletion when the intermediate homocysteine is provided (1,2). Most normal cells experience a small and slow decrease in proliferation under methionine depletion. Addition of homocysteine to the culture media completely rescues proliferation. Cancer cells, on the other hand, die within 48-72h after the initiation of methionine depletion, and rescue by homocysteine is modest (1). Exceptions exist to this rule. Activated T cells are not rescued by homocysteine (3). Conversely, rare cancer cell lines, such as the human melanoma MeWo, can be rescued nearly completely with homocysteine (3). *In vivo*, enzymatic and dietary methionine depletion reduces tumor growth and synergizes with many cancer therapies (5–9). At the same time, dietary methionine restriction is associated with improvements in glucose and lipid regulation, and in a general increase in lifespan (10). However, the molecular origin of the difference in response to methionine depletion between methionine independent and methionine dependent cells remains unknown.

Methionine is an essential amino acid fulfilling several crucial metabolic roles. In addition to protein synthesis, methionine is the precursor for the universal methyl donor S-adenosylmethionine. S-adenosyl methionine donates methyl groups to several acceptor molecules such as DNA, RNA, proteins (including, but not restricted to, histones), creatine, and phospholipids. The resulting demethylated molecule, S-adenosylhomocysteine, yields homocysteine which can then be either enter the remethylation pathway to methionine or the transulfuration pathway to the antioxidant glutathione. Alternatively, S-adenylsylmethionine can be decarboxylated and enter the methionine salvage pathway, resulting in the production of polyamines.

While methionine synthase expression is similar between methionine dependent and independent cell lines, there may be differences in the rates of methionine remethylation and transmethylation (11). Treatment with an inhibitor of DNA methyltransferase was also found to cause a reversion from dependence to independence (11). Comparison of the methionine independent MeWo cell line with its more aggressive, methionine dependent variant MeWo-LC1 identified that hypermethylation of the promoter region of the vitamin B_12_ chaperone MMACHC explained the divergence in methionine phenotype (4). However, this finding was limited to this single cell line. The use of methyl groups is also different between methionine dependent and independent cells. The methyl group from methionine is used primarily for the synthesis of S-adenosylmethionine and methylated proteins in methionine independent cells (12). Methionine independent cells, on the other hand, primarily use this methyl group for the synthesis of creatine and phosphatidylcholine (12).

In this study, we sought to understand the many ramifications of the replacement of methionine with homocysteine in cell culture media. We compared the methylome and proteome of two human melanoma cell lines. One, A101D, is a prototypical cancer cell line displaying methionine dependence. The second, MeWo, has long been described as methionine independent, i.e. rescued by homocysteine. We identified a general decrease in promoter DNA methylation, with changes in the cellular transcriptome and proteome. Despite apparently sensing equally the shift from methionine to homocysteine, the effector response was noted to be much larger in methionine dependent than in methionine independent cells. Our analysis reveals the importance of cell cycle control, DNA damage repair, translation, nutrient sensing, oxidative stress, and tight junctions in the cellular response to methionine stress in melanoma.

## Methods

### Cell culture

A101D and MeWo cells were obtained from ATCC. Cells were cultured in high glucose, no glutamine, no methionine, no cystine DMEM (ThermoFisher Scientific, Waltham MA) supplemented with 5% dialyzed serum (BioTechne, Minneapolis, MN), 100 IU penicillin and streptomycin (ThermoFisher), 4 mM L-glutamine (ThermoFisher) and 1 mM sodium pyruvate (ThermoFisher). L-cystine (Millipore-Sigma, Burlington, MA) was resuspended in PBS with NaOH added until complete solubilization, and added to the cell media at a final concentration of 150 μM. For control medium, L-methionine (Millipore-Sigma) was resuspended in PBS and added to the cell media at a final concentration of 200 μM. For methionine-free, homocysteine containing media, L-homocysteine (Chem-impex, Wood Dale, IL) was resuspended in 1M HCl in PBS and added to the cell media at a final concentration of 200 μM.

### Cell cycle

A101D and MeWo cells were cultured in either methionine or homocysteine media for 24 hours or 48 hours. Cells were fixed in ethanol, washed, and treated with ribonuclease. Finally, cells were stained with DAPI and analyzed for cell cycle by fluorescence activated sorting on a BD LSRFortessa instrument (Beckman Coulter, Brea, CA).

### DNA methylation

A101D and MeWo cells were grown in either control media or homocysteine media for 24 hours. DNA and RNA was extracted using the AllPrep kit following the manufacturer’s instructions (Qiagen, Germantown, MD). After Qubit fluometric quantification (ThermoFisher), methylation was analyzed using the Infinium MethylationEPIC array (Illumina, San Diego, CA) at the Genomics Core of the University of Arkansas for Medical Sciences (UAMS). Data analysis was performed at the UAMS Bioinformatics Core. Methylation was expressed as the absolute difference between cells grown in methionine and cells grown in homocysteine. The data is available in the Gene Expression Omnibus database under accession number GSE225946.

### RNA-Seq

Cells were culture for 24 hours in media containing 200 μM methionine or 200 μM homocysteine. RNA was extracted with a commercially available kit (RNeasy, Qiagen). After Qubit fluometric quantification (ThermoFisher), the NGS library was prepared with the Truseq Stranded Total RNA (Illumina, San Diego, CA) and ran on a HiSeq300 instrument at the UAMS Genomics Core. Data analysis was performed at the UAMS Bioinformatics Core. The data is available in the Gene Expression Omnibus database under accession number GSE225945.

### Cell cycle

Cell cycle was assessed by flow cytometry. Cells were culture for 24 and 48 hours in media containing 200 μM methionine or 200 μM homocysteine. The cells were harvested then fixed in 70% ethanol. The day of the assay, the cells were rinsed in PBS and incubated in a solution containing 100 μg/mL RNAse A and 1 μg/mL DAPI. Analysis was performed at the UAMS flow cytometry core on a BD LSR Fortessa (BD, Franklin Lakes, NJ) instrument and using the FlowJo software (BD).

### Proteomics

Purified proteins were reduced, alkylated, and digested using filter-aided sample preparation (13). Tryptic peptides were labeled using a tandem mass tag 10-plex isobaric label reagent set (ThermoFisher) following the manufacturer’s instructions. Labeled peptides will be separated into 36 fractions, and then consolidated into 12 super-fractions. Each super-fraction was further separated by reverse phase. Peptides were eluted and ionized by electrospray followed by mass spectrometric analysis on an Orbitrap Fusion Lumos mass spectrometer (ThermoFisher). MS data were acquired using the FTMS analyzer and MS/MS data using the ion trap analyzer. Up to 10 MS/MS precursors were selected for HCD activation, followed by acquisition of MS3 reporter ion data. Proteins were identified and reporter ions quantified by searching the UniprotKB database using MaxQuant (Max Planck Institute) with a parent ion tolerance of 3 ppm, a fragment ion tolerance of 0.5 Da, and a reporter ion tolerance of 0.001 Da. Protein identifications were accepted if they can be established <1.0% false discovery and contained at least 2 identified peptides. Protein probabilities were assigned by the Protein Prophet algorithm (14). TMT MS3 reporter ion intensity values were log2 transformed and missing values imputed by a normal distribution for each sample using Perseus (Max Planck Institute). TMT batch effects were removed using ComBat (15). Statistical analysis was performed using Linear Models for Microarray Data with empirical Bayes smoothing to the standard errors (16). Proteins with an FDR adjusted p-value <0.05 were considered to be significant. The data is available in the ProteomeXchange database under the ID PXD040655 and MassIVE MSV000091423.

### IPA analysis

Significant proteins (FDR adjusted p-value <0.05) were utilized to identify important protein networks and pathways using Qiagen’s Ingenuity Pathway Analysis (17). For A101D RNA-Seq analysis, we limited the dataset to fold change >1.5X.

## Results

### Homocysteine induces hypomethylation in both MeWo and A101D cells

Methionine is an important methyl donor in cells and we therefore investigated the DNA methylation patterns in methionine dependent and independent human melanoma cells cultured with either methionine or homocysteine. We chose a time point of 24 hours to measure cells prior to cell death. Beta values were significantly decreased in homocysteine versus methionine for both cell lines, indicating a net decrease in methylation levels (Fig. 1A). Interestingly, the beta value was decreased to a greater extent in MeWo than in A101D, despite the cell cycle phenotype being significantly milder. Accordingly, a total of 252 probes were significantly differentially methylated between methionine and homocysteine in the methionine dependent A101D, compared to 1145 in MeWo (Supplementary data). All significant sites in both cell lines were hypomethylated in homocysteine compared to methionine. IPA analysis suggested that Sonic Hedgehog signaling was the main canonical pathway dysregulated in A101D, and calcium signaling in MeWo (Fig 1B). However, the relatively large p values associated with these keywords suggest a low-level enrichment.

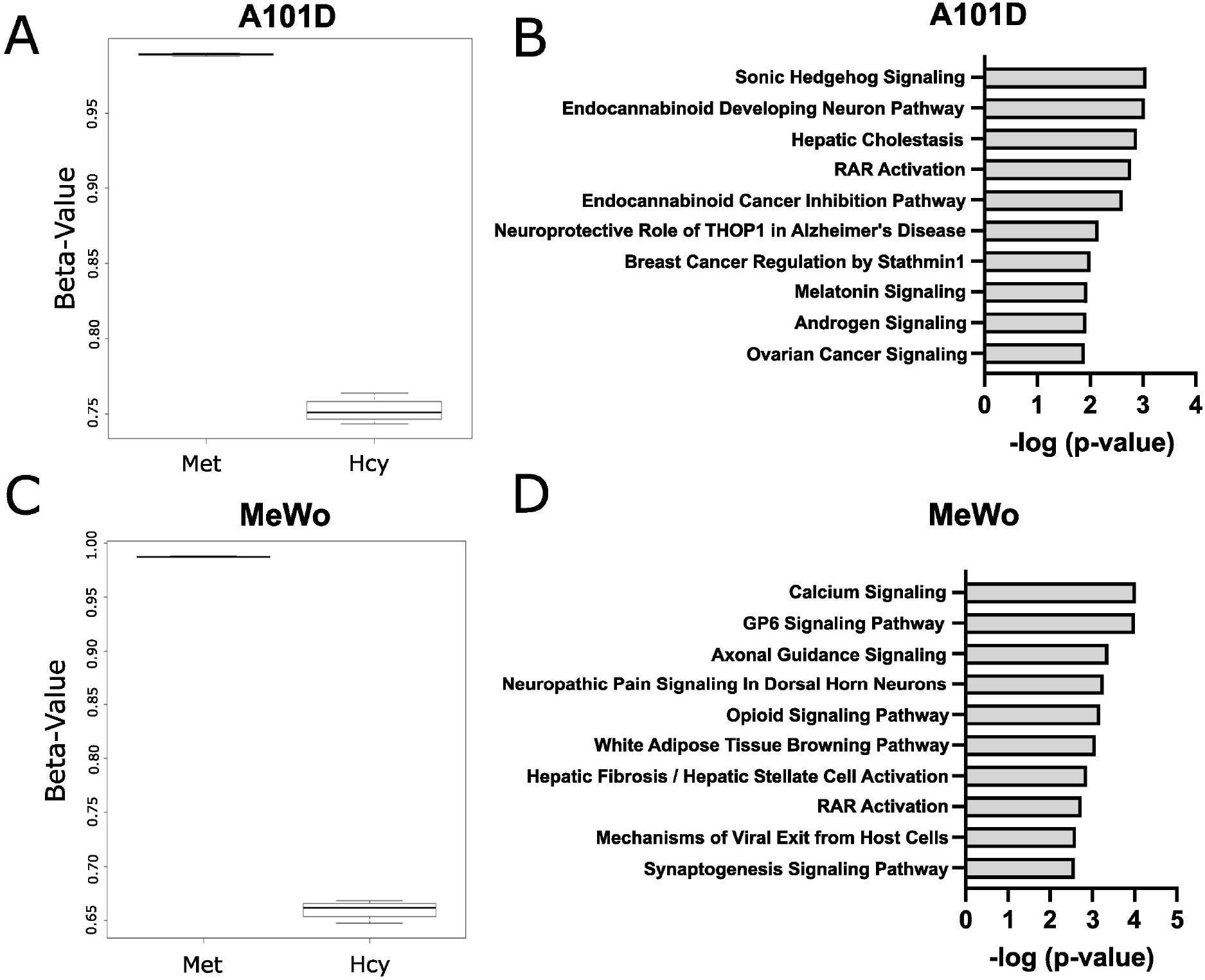

### Homocysteine is associated with a stronger transcriptomic signature in methionine-dependent A101D than MeWo

In order to investigate further the signaling events suggested by the methylation array, we analyzed the transcriptome of cells growing in either methionine or homocysteine. As opposed to DNA methylation, the replacement of methionine with homocysteine induced much greater transcriptome-level changes in the methionine dependent cell line than in the methionine independent cell line. In the methionine-dependent A101D, over 12,000 genes were significantly altered, as defined by an adjusted p value < 0.05. This included 2580 genes altered with a change of more than 2X (1545 up and 1035 down)(Supplementary data).

In comparison, only 3843 genes were significantly altered in MeWo under the same culture conditions. Of these, only 110 by more than 2X (58 up and 52 down). To control for any experimental variable, we looked at *MAT2A*, the gene encoding methionine adenosyltransferase 2A. MAT2A catalyzes the conversion of methionine to S-adenosylmethionine and responds strongly to methionine stress. We found that MAT2A was increased 4.5X in A101D and 3.5X in MeWo following the switch to homocysteine. This indicated that the absence of methionine was sensed nearly equally in both cell lines. MAT2A was in fact the gene with the smallest adjusted p value in MeWo (6.35×10^-33^).

We then analyzed pathway enrichment with IPA. Taking advantage of the large number of significantly altered target in A101D, we refined our analysis by separating upregulated and downregulated genes. In that methionine-dependent cell line, there was an enrichment for pathways related to the immune system among upregulated genes (Fig 2A). The analysis pointed to cytokines and pattern recognition receptors. This was somewhat unexpected but is in good agreement with new results from our laboratory showing that methionine restriction activates immune pathways in cancer (BIORXIV/2023/535695). This was also reflected in the upstream analysis, where the top hits were the immune-related factors TNF, IFNG, and IL1B.

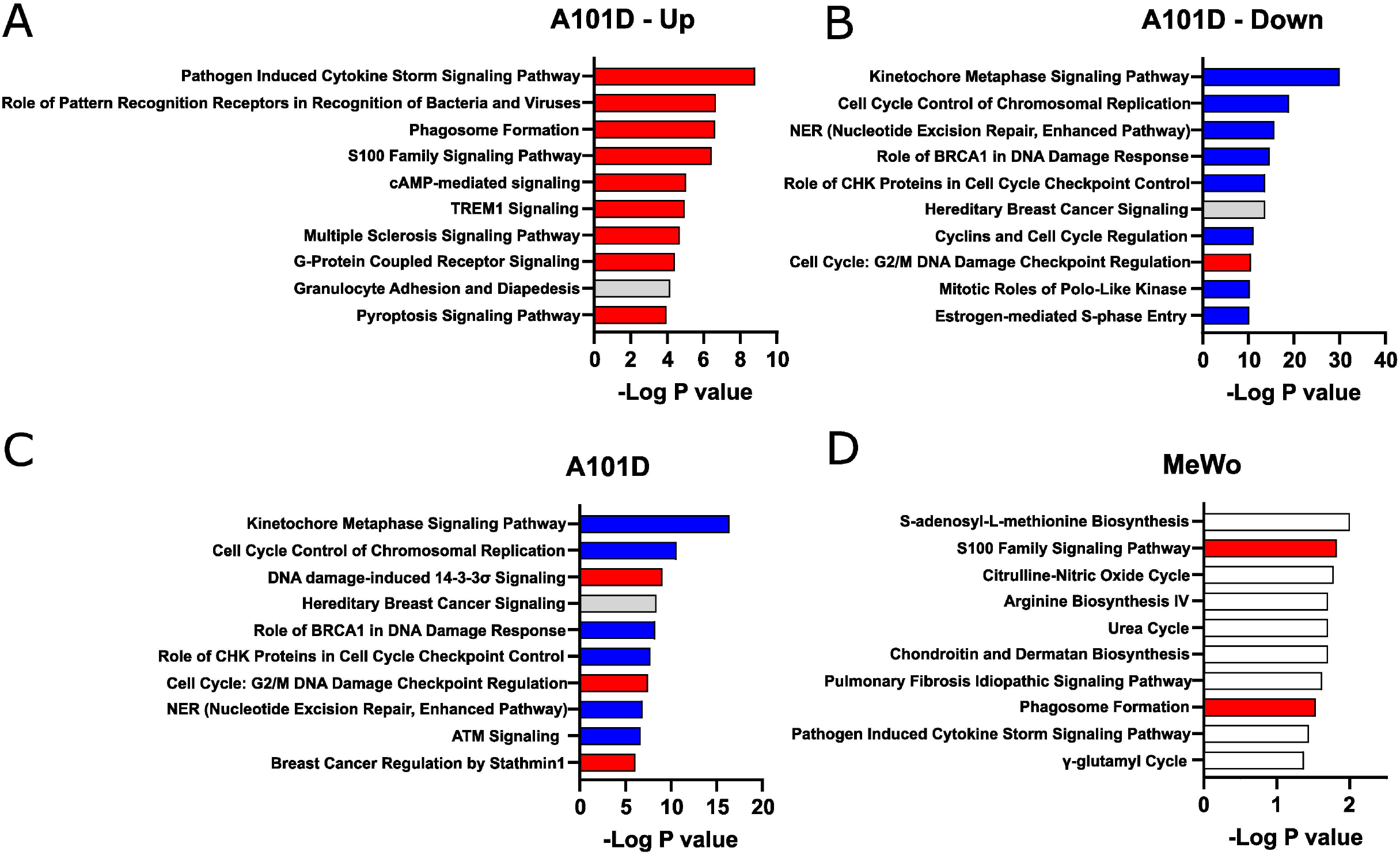

Pathway enrichment was noticeably greater in genes downregulated in A101D (Fig 2B) than in any other condition presented in Figure 2, as reflected by the different scale on the X axis. In methionine-dependent A101D, pathways related to the cell cycle included “kinetochore metaphase signaling pathway” and “cell cycle control of chromosomal replication”. Genes supporting this pathway included centromere proteins and various components of the KMN network. Pathways related to DNA damage included “NER (nucleotide excision repair, enhanced pathway)”, and “role of BRCA1 in DNA damage response”. Genes supporting this pathway included genes encoding FANC proteins, *BRCA1* and BRCA1 complex genes, *RAD51* and *TP53*. The upstream analysis identified L-asparaginase as the top hit. This further confirms that the changes related to methionine dependence are mostly related to its role as a protein building amino acids rather than as a methyl donor.

In methionine-independent MeWo, the limited number of genes altered by more than 2X led to pathways with lower confidence values (Fig 2CD). The pathway “S-adenosyl-L-methionine biosynthesis” was supported solely by *MAT2A*. “S100 family signaling pathway”, however, was supported by 7 genes and reflected the same keyword found in A101D. We decreased the threshold to 1.5X change without observing significant differences (data not shown). Other genes downregulated in MeWo included argininosuccinate synthase 1, which supported the keywords “citrulline-nitric oxide cycle”, “arginine biosynthesis IV”, and “urea cycle”.

One of the largest fold change increases in A101D was the gene *FGF21* (38X, adj p value 4×10^-4^). FGF21 regulates the metabolic adaptations resulting from protein restriction in general (18,19), and methionine restriction in particular (20). FGF21 is a hepatic protein that plays an essential role in increasing insulin sensitivity, increasing energy expenditure, and activating thermogenesis in animals fed a methionine-restricted diet (20). Its role in cancer is less well defined but is thought to involve the SIRT1/PI3K/AKT axis (21). Another growth factor, growth differentiation factor 15 (GDF15), was also among the most highly upregulated genes in A101D. GDF15 has important metabolic functions and has been reported to be upregulated during mitochondrial dysfunction (22,23). *FGF21* and *GDF15* were not significantly upregulated in MeWo.

### Homocysteine affects the cell cycle in methionine dependent A101D but not methionine independent MeWo

To confirm the effect of methionine versus homocysteine on the cell cycle seen with the transcriptomics in A101D human melanoma cells, we performed a cell cycle analysis after 24 and 48 hours. After 24 hours, there was an increase in cells in the G1 phase and a decrease in cells in the S and G2-M phases in A101D. MeWo cells, on the other hand, had a decrease in G1 cells and an increase in G2-M cells (Fig 3AB). After 48 hours, there was a large increase in SubG1 cells that was specific to A101D (Fig 3C). This indicates the presence of fragmented DNA, in agreement with the decrease in DNA repair keywords in the transcriptomics. There was a decrease in cells in G1, S, and G2-M. In MeWo cells, there was still an increase in G2-M cells at 48h, with a decrease in cells in S phase (Fig 3D). This suggests that A101D methionine dependent cells first reduce their proliferation, then undergo apoptosis (24). Cell division may be briefly promoted by homocysteine in MeWo methionine independent cells.

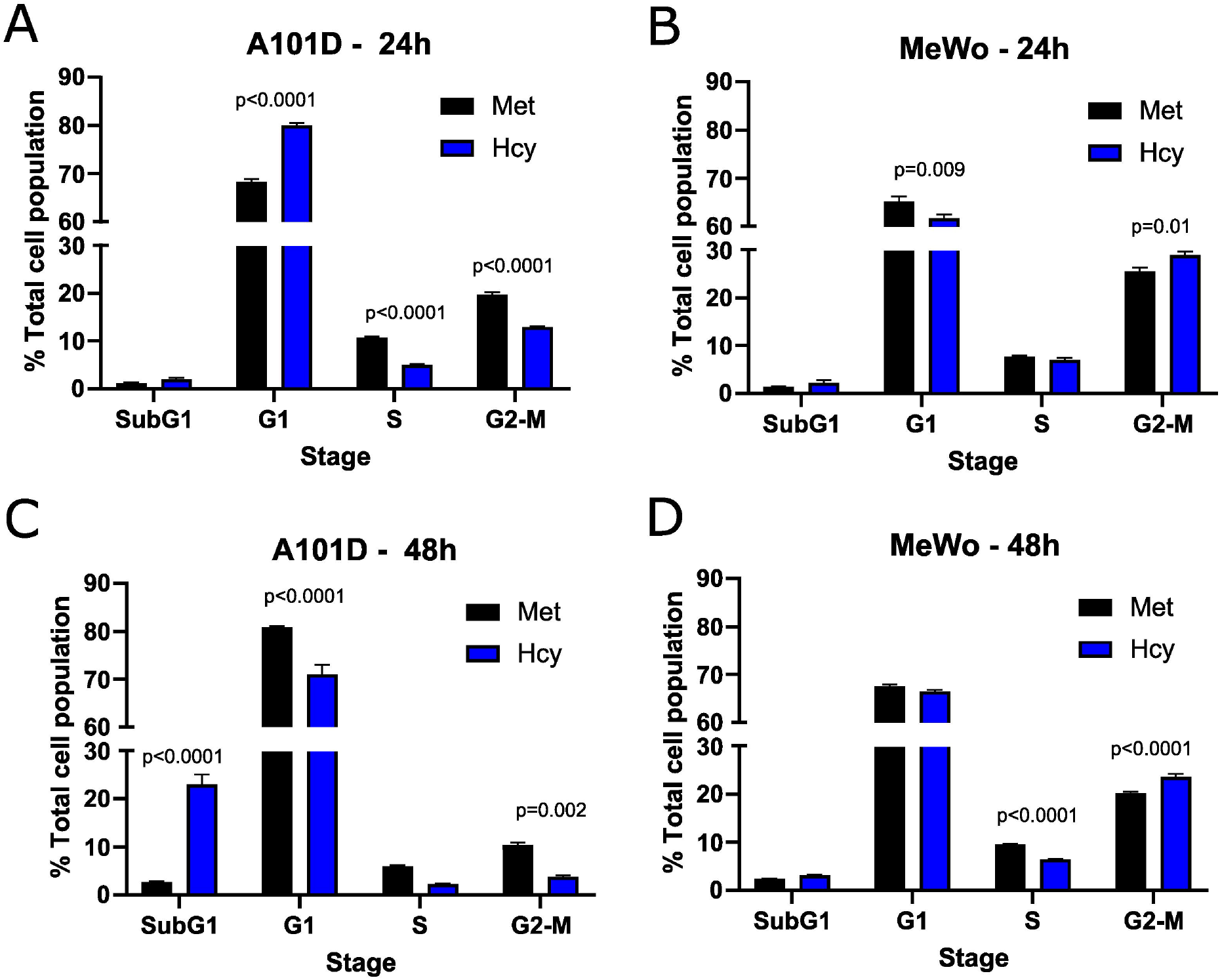

### Proteomics provide additional insight into methionine dependence

To gain additional information about the differences between methionine dependent and independent, we performed an analysis of the cellular proteome of A101D and MeWo cells in methionine versus homocysteine-containing media. This type of analysis provides further information about translational and post-translational changes in the cell. As for the transcriptomics data, the changes were more pronounced in A101D than MeWo (Fig 4 and Supplementary data). We first noticed that there were fewer significant changes in the proteomics analysis then in the transcriptomic analysis. Few displayed changes larger than 2X (4 proteins in A101D and 3 proteins in MeWo). Using a lower threshold of 1.25X change, there were 345 significantly altered proteins in A101D (142 downregulated and 203 upregulated) and only 115 proteins in MeWo (6 downregulated and 109 upregulated). We proceeded with the IPA analysis with the 1.25X threshold. There was a sufficient number of proteins in A101D to split the analysis between upregulated and downregulated proteins, but not in MeWo. Therefore, we are showing upregulated proteins (Fig. 4A), downregulated proteins (Fig. 4B), and combined (Fig. 4C) for A101D, and combined for MeWo (Fig. 4D).

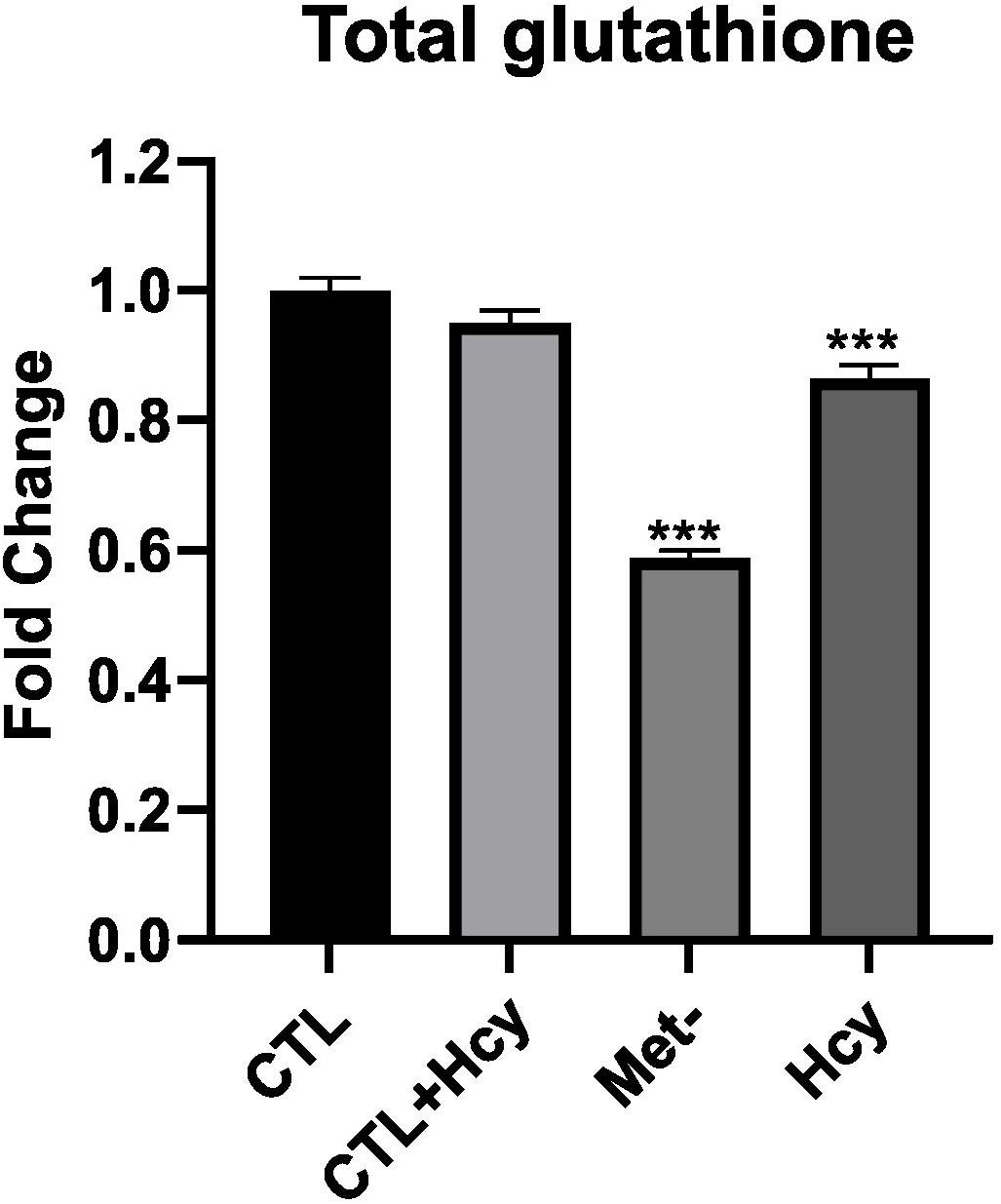

Based on our transcriptomics data, we looked more closely at the abundance of the protein MAT2A. As suggested by the transcriptomic analysis, we found that the protein level of MAT2A was increased for both A101D and MeWo (1.6X and 1.5X, respectively). Along with the gene expression data, this confirms previous reports in methionine dependent cells (25) and expands this observation to methionine independent cells.

Among upregulated proteins in A101D, the top pathway was “NRF2-mediated oxidative stress response”, with contributing increased proteins such as glutathione-S-transferases, PRDX1, SOD1/2, and TNX. This is in good agreement with previous reports (24,26). The glutathione-S-transferases also contributed to all other pathways in the top 10, except “sirtuin signaling pathway” and “glycolysis I”. We had previously shown that total glutathione is decreased by about 40% in A101D cells grown in homocysteine compared to cells grown in methionine (24). MeWo cells, on the other hand, show only a 14% decrease in total glutathione (Fig 5). Notably, the proteomic analysis did not indicate any enrichment for immune pathways, contrarily to the transcriptomic analysis. The growth factor FGF21 was not detected in our proteomics analysis. However, GDF15 was the most upregulated protein in A101D (3.7X), mirroring the gene expression data. The protein GDF15 was also upregulated 1.4X in MeWo.

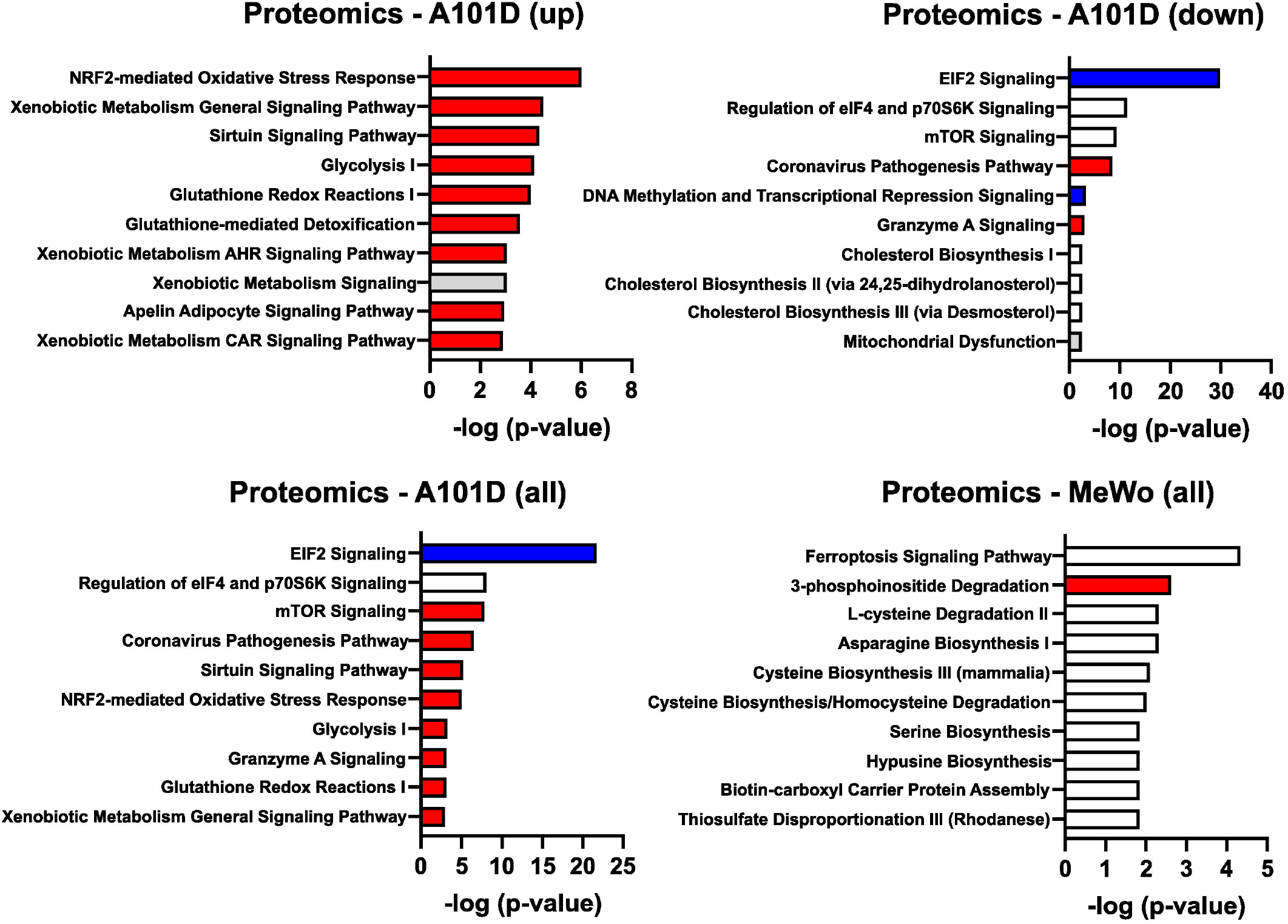

Despite the larger number of upregulated versus downregulated proteins in A101D, it did not yield the largest pathway enrichment. It was rather found among downregulated proteins in A101D for the pathway “EIF2 signaling”. This analysis indicates a decrease in translation, caused in part by a decrease in the abundance of over 25 different ribosomal proteins. In contrast, no ribosomal protein was significantly altered by more than 25% in the methionine-independent MeWo. This is in good agreement with the data we published previously showing that proliferation was severely reduced in A101D cells grown in homocysteine compared to methionine, while being unchanged in MeWo (24). Analysis of the transcriptomics dataset showed similar changes, but it did not add to a significant pathway enrichment. Other pathways enriched in the downregulated proteins included “regulation of eIF4 and p70S6K signaling” and the related “mTOR signaling”, which responds to nutrient abundance and is regulated by the level of S-adenosylmethionine in the cell (27). The proteins contributing to this pathway were for the most part a subset of ribosomal proteins contributing to “EIF2 signaling”. The same was true for the keyword “coronavirus pathogenesis pathway”. These four top pathways in the downregulated proteins also formed the top 4 pathways for the combined upregulated and downregulated proteins (Fig. 3C).

The single pathway suggesting an involvement of DNA methylation in methionine dependence was found among downregulated proteins in A101D. Components of the NuRD and Sin3 complexes of histone and DNA modifications contributed to the pathway “DNA methylation and transcriptional repression signaling”. Interestingly, the analysis of downregulated proteins in A101D cells grown in homocysteine also produced an enrichment for pathways related to cholesterol. Methylsterol monooxygenase 1 and squalene epoxidase were both significantly decreased. This echoed the gene expression findings in MeWo, indicating a possible involvement of cholesterol and cholesterol-related molecules in the response to methionine stress both in methionine dependent and independent cells.

The top pathway enriched in MeWo cells was “ferroptosis signaling pathway”. It reflected an increase in proteins that have been linked to lipid peroxidation, such as the amino acid transporter SLC1A5, ferritin heavy chain1, cystathionine gamma-lyase, ADP ribosylation factor 5, and the transcription factor SP1. Both ferritin light chain and ferritin heavy chain were increased in A101D (1.7X and 1.3X, respectively). We have previously reported on the ferroptosis-like gene expression signature in methionine-restricted cells. We and others have shown that this gene signature is not accompanied by an increase in either lipid peroxidation nor death by ferroptosis, contrarily to cysteine depletion (24,28). Interestingly, the transferrin receptor was decreased both in A101D and in MeWo (0.6X and 0.8X, respectively). There was also an enrichment in the keyword “3-phosphoinositide degradation”, supported by an increase in MTMR1, MTMR14, NUDT3, PIP4P2, and SSH3. Finally, individual or pairs of genes contributed to some amino acid biosynthesis and degradation pathways.

Finally, two cobalamin-related proteins were identified in the proteomics screen. Cobalamin is the cofactor for the cytoplasmic methionine synthase reaction, as well as the mitochondrial methylmalonyl-CoA mutase reaction. MMAA translocates cobalamin from the cytosol into the mitochondrion and was increased 1.7X in MeWo. The lysosomal cobalamin transporter ABCD4 was increased 1.8X in A101D.

## Discussion

Cancer cells are generally more resistant to harsh environmental conditions than normal cells. However, they also display specific vulnerabilities. Methionine dependence in cancer cells is one of such vulnerabilities, described over 45 years ago. We attempted to characterize the molecular bases behind the inability of most cancer cells to utilize homocysteine to sustain growth and metabolism.

A single methyl group differentiates methionine from homocysteine. We therefore looked at differences in DNA methylation between A101D and MeWo. DNA hypomethylation was a hallmark in both cell lines, but interestingly, the effect was more profound in MeWo cells. Continued proliferation in these cells in methionine-free, homocysteine-containing media may lead to a dilution effect compared to non-proliferating A101D cells. Although some signaling pathways were significantly enriched, both positive and negative regulators were hypomethylated, leading to an uncertain outcome on cellular function. Others have shown alterations in histone modification in connection with methionine metabolism (29–32). Our data suggest that these modifications may be more relevant to the control of gene expression following methionine depletion then DNA methylation.

To gain more insight into differences in metabolism between methionine dependent and methionine independent melanoma cells, we turned our attention to the analysis of the transcriptome and proteome. This revealed very clearly that methionine dependent cells show much greater changes in response to the change from methionine to homocysteine compared to methionine independent cells. In this experiment, the transcriptomics analysis performed better at identifying alterations in the cell cycle in methionine dependent cells. The impact on the cell cycle was readily validated by flow cytometry. As expected, proliferation was decreased at both 24 and 48h in A101D cells, in contrast to a slight increase in cells undergoing mitosis in MeWo. An increase in cells in the subG1 phase in A101D cells after 48h suggested an apoptotic mechanism of cell death in methionine dependent cells grown in methionine-free, homocysteine-containing media. Transcriptomics also put more emphasis on DNA damage.

A novel finding revealed during this analysis was the enrichment of immune pathways in methionine dependence, particularly cytokines and pattern recognition receptors. Interestingly, lymphocytes are the only non-cancer cell type reported to be methionine dependent (3). More recently, methionine uptake has been shown to crucial to the activation and proliferation of T cells (33–35). However, dietary methionine restriction was also shown to delay age-related changes in T cell subsets in mice, drawing a more complex picture of the immune effects of the treatment (36). Finally, reducing methionine in the diet has been reported to promote anti-tumor immunity (37). Our own results indicate that methionine restriction promotes the cGAS-STING pathway (submitted) in a similar fashion as arginine restriction (38). None of the immune-related change in the methionine dependent A101D were identified in the methionine independent MeWo. This indicates that the immune changes produced by alterations in methionine may be confined to methionine dependent cancer cells and T cells.

In this experiment, proteomics provided a different picture of the response to methionine stress. The proteomics analysis in MeWo revealed a limited number of significant changes and revealed fewer insights. In A101D however, the analysis revealed an increase in oxidative stress. This was specifically driven by an increase in proteins involved in glutathione synthesis. In cells with an active transulfuration pathway such as hepatocytes and cancer cells, methionine serves as a precursor for the main cellular antioxidant glutathione. The decrease in glutathione measured in the cells suggests that the activation of this pathway results from a decrease in products.

Another theme that was uniquely contributed by the proteomic analysis was a decrease in translation, driven by a decrease in ribosomal proteins. Processes such as ribophagy have been described that would fit a selective degradation of ribosomal proteins. However, the transcript-level downregulation of the ribosome receptor for ribophagy NUFIP1 (39) is downregulated (Log fold change = −0.31, −Log p value = 7.9) in these cells at 24 hours argues against this possibility. Given that translation lags behind transcription, there may rather be a rapid and transient decrease in the expression of the genes encoding these proteins that is resolved by our time point of 24 hours, leaving only protein levels decreased.

In summary, we found little evidence for the role of DNA methylation patterns in methionine dependence. We did however find strong evidence that the decrease in methionine is equally sensed by both methionine dependent and independent cells, leading to a reliable increase in MAT2A gene ad protein expression in both methionine dependent and independent cell lines. This is different to what was previously published in glioma cells (40), but in good agreement with other results in breast cancer (25). The changes engendered in the methionine independent MeWo cell line were generally also found in the methionine dependent A101D. However, additional layers of changes were also observed in the latter, specifically oxidative stress, translation, and cell cycle control.

## Supporting information

Supplementary Data - Methylation

Supplementary Data - RNA-Seq

Supplementary Data - Proteomics

## Acknowledgements

The authors would like to thank Drs. Giulia Baldini and Nükhet Aykin-Burns for guidance. Lisa Orr and Owen Stephens expertly processed samples at the National Resource for Quantitative Proteomics and the UAMS Genomics core. Kathy Bush provided emergency childcare during the COVID-19 crisis. This work was supported by the Winthrop P. Rockefeller Cancer Institute (WPRCI), the Arkansas Tobacco Settlement Commission, the National Center for Advancing Translational Sciences grant number 1KL2TR003108-01, and NIGMS grant P20 GM139768-01. The National Resource for Quantitative Proteomics is supported by the National Institute of General Medical Sciences (NIGMS) grant R24GM137786. The UAMS Bioinformatics core is supported by NIH/NIGMS grant P20GM121293 and WPRCI.

## Notes

https://www.ncbi.nlm.nih.gov/geo/

https://proteomecentral.proteomexchange.org/cgi/GetDataset

### Competing Interest Statement

The authors have declared no competing interest.

